# Facile semi-automated forensic body fluid identification by multiplex solution hybridization of NanoString^®^ barcode probes to specific mRNA targets

**DOI:** 10.1101/007898

**Authors:** Patrick Danaher, Robin Lynn White, Erin K. Hanson, Jack Ballantyne

**Author notes:** Correspondence: Jack Ballantyne, Ph.D. Professor Department of Chemistry Associate Director (Research) National Center for Forensic Science University of Central Florida PO Box 162367 Orlando, FL 32816-2367 Voice.

## Abstract

A DNA profile from the perpetrator does not reveal, *per se*, the circumstances by which it was transferred. Body fluid identification by mRNA profiling may allow extraction of contextual ‘activity level’ information from forensic samples. Here we describe the development of a prototype multiplex digital gene expression (DGE) method for forensic body fluid/tissue identification based upon solution hybridization of color-coded NanoString^®^ probes to 23 mRNA targets. The method identifies peripheral blood, semen, saliva, vaginal secretions, menstrual blood and skin. We showed that a simple 5 minute room temperature cellular lysis protocol gave equivalent results to standard RNA isolation from the same source material, greatly enhancing the ease-of-use of this method in forensic sample processing.

We first describe a model for gene expression in a sample from a single body fluid and then extend that model to mixtures of body fluids. We then describe calculation of maximum likelihood estimates (MLEs) of body fluid quantities in a sample, and we describe the use of likelihood ratios to test for the presence of each body fluid in a sample. Known single source samples of blood, semen, vaginal secretions, menstrual blood and skin all demonstrated the expected tissue-specific gene expression for at least two of the chosen biomarkers. Saliva samples were more problematic, with their previously identified characteristic genes exhibiting poor specificity. Nonetheless the most specific saliva biomarker, HTN3, was expressed at a higher level in saliva than in any of the other tissues.

Crucially, our algorithm produced zero false positives across this study’s 89 unique samples. As a preliminary indication of the ability of the method to discern admixtures of body fluids, five mixtures were prepared. The identities of the component fluids were evident from the gene expression profiles of four of the five mixtures. Further optimization of the biomarker ‘CodeSet’ will be required before it can be used in casework, particularly with respect to increasing the signal-to-noise ratio of the saliva biomarkers. With suitable modifications, this simplified protocol with minimal hands on requirement should facilitate routine use of mRNA profiling in casework laboratories.

## 1. Introduction

Genetic identification of the donor of transferred biological traces deposited at the crime scene or on a person using STR analysis is now routine practice worldwide [1]. This represents potentially crucial ‘source level’ information for investigators [2]. A DNA profile from the perpetrator does not however reveal the circumstances by which it was transferred. This contextual information (sometimes known as the ‘activity level’ in Cook and Evett’s classic 1998 paper [2]) is important for casework investigations because the deposition of the perpetrator’s biological material requires some behavioral activity that results in its transfer from the body. The consequences of different modes of transfer of the DNA profile may dramatically affect the investigation and prosecution of the crime. For example a DNA profile from a victim that originates from skin versus the same DNA profile that originates from vaginal secretions may support social or sexual contact respectively. Thus tissue/body fluid sourcing of the DNA profile should be an important concern for, and service from, forensic genetics practitioners who are integral to the investigative team. The problem is that, up until the recent past, it was not possible to definitively identify many of the important body fluids of interest (e.g. vaginal secretions, saliva, and menstrual blood).

In order to overcome the limitations of currently used classical body fluid identification methods, the use of messenger RNA (mRNA) profiling, as described by Juusola & Ballantyne [3], was proposed to supplant conventional methods for body fluid identification. Terminally differentiated cells, whether they comprise blood monocytes or lymphocytes, ejaculated spermatozoa, epithelial cells lining the oral cavity or epidermal cells from the skin become such during a developmentally regulated program in which certain genes are turned off (i.e. transcriptionally silent) and turned on (i.e. are actively transcribed and translated into protein) [4]. Thus, a pattern of gene expression is produced that is unique to each cell type in both the presence and the relative abundance of specific mRNAs [4]. The type and abundance of mRNAs, if determined, would then permit a definitive identification of the body fluid or tissue origin of forensic samples. This is the basis for mRNA profiling for body fluid identification. RNA profiling now offers the ability to identify all forensically relevant biological fluids using methods compatible with the current DNA analysis pipeline [5,6]. Despite the identification of numerous body fluid specific candidates there is some reluctance to utilize RNA profiling assays in the forensic community due to concerns over the perceived instability of RNA in biological samples. However, several studies have been conducted in order to assess the stability of RNA in dried forensic stains [7–10]. These have demonstrated that RNA of sufficient quantity and quality for analysis can be recovered from aged and environmentally compromised forensic samples [7–10]. The effective stability (i.e. ‘recoverability’) of mRNA in aged and compromised samples is not dissimilar to that of DNA and provides support to the use of mRNA profiling assays in forensic casework (Ballantyne, unpublished observations). The recently published EDNAP collaborative exercises on mRNA profiling for body fluid identification further demonstrate a significant interest in mRNA profiling by the forensic community in Europe and around the world as well as the ease in which this technology can be implemented into forensic casework laboratories [11–15]. Collectively, these studies demonstrate an interest in the use of mRNA profiling in forensic casework and its suitability of use with forensic samples and therefore warrant continued evaluation and development. Other classes of RNA also exist in the cell and one in particular, microRNA (miRNA), has been investigated for potential forensic use since the short size of the molecule (∼21–25 bases) makes it an attractive option for analyzing degraded specimens [16–22]. The field of forensic miRNA profiling, although promising, is less mature in terms of there being an international consensus on the identity and specificity of the best body fluid specific miRNA targets. Other non-RNA methods for body fluid identification have been recently investigated including the use of epigenetic [23–29] and proteomic [30–32] biomarkers. Although exhibiting some promise, epigenetic markers have not been identified for all of the important common body fluids and tissues such as vaginal secretions and skin. Proteomic markers suffer from a lack of demonstrated reproducibility studies among different laboratories, and paucity of peer reviewed reports demonstrating their forensic validity.

Gene expression differences are quantitative in nature meaning that a particular biomarker may be expressed in a particular cell type at low, intermediate or high levels. Even when it is not generally regarded as being expressed in a particular cell type it may exhibit basal level (or ‘leaky’) transcription with a few molecules present per cell. Thus far there have been three main methods developed for mRNA profiling of forensic samples: capillary electrophoresis (CE)-based analysis [5–7,33–36], quantitative RT-PCR (qRT-PCR) [7,37–39] and, more recently, high resolution melt (HRM) analysis [40]. Due to its facile multiplex capabilities and routine use in DNA profiling, CE-based analysis has been the platform of choice for casework mRNA assays [5,6]. However post–PCR CE peak heights/areas are, at best, semi-quantitative in nature with respect to biomarker expression levels. Similarly, HRM signal amplitude does not appear to correlate precisely with RNA input [40]. Although qRT-PCR permits quantitation of biomarker targets, its low multiplex capability (typically 3-4 targets maximum compared to >20 for CE) appears to have limited its use.

In contrast to the aforementioned, digital gene expression (DGE) methods precisely count the number of individual transcripts in a sample [41] which facilitates the use of advanced statistical methods to better evaluate and interpret the experimental data. This facility would be expected to be of significant benefit when analyzing body fluid mixtures that are commonly encountered in forensic analysis. Deep sequencing of the transcriptome using next generation sequencing (NGS) technologies is capable of directly identifying and quantifying (by counting) all mRNA transcripts in a sample, a DGE technique known as RNA sequencing (RNA-Seq) [42]. RNA-Seq has been spectacularly successful in advancing our knowledge of cell-type-specific gene expression including transcript quantification and elucidation of their sequence diversity [42]. Although NGS heralds a new era of forensic genomics, impediments to its routine implementation in body fluid RNA analysis include its high cost of reagents and time-consuming, complex analysis. In this work we sought an alternative DGE method to NGS that is simpler and requires minimal hands-on experimentation. Here we describe the development of a prototype multiplex DGE method for forensic body fluid identification based upon solution hybridization of color-coded NanoString^®^ probes [43] to 23 tissue/body fluid specific and 10 housekeeping gene mRNA targets present in forensic type samples. Concomitantly, to facilitate routine use, we also devised a simple 5 minute room temperature cellular lysis protocol as an alternative to standard RNA isolation for forensic sample processing.

## 2. Methods

### 2.1 Body fluid samples

Body fluids were collected from volunteers using procedures approved by the University’s Institutional Review Board. Informed written consent was obtained from each donor. Blood samples were collected by venipuncture into vacutainers (K3-EDTA preservative) and 50 μl aliquots were placed onto cotton cloth and dried at room temperature. Freshly ejaculated semen was provided in sealed plastic tubes and stored frozen. After thawing, the semen was absorbed onto sterile cotton swabs and allowed to dry. Buccal samples (saliva) were collected from donors using sterile swabs by swabbing the inside of the donor’s mouth. Semen-free vaginal secretions and menstrual blood were collected using sterile cotton swabs. Admixed body fluid samples were created by combining ½ of a 50 μl stain or single cotton swab from each body fluid. Environmental samples were prepared by exposing body fluid samples to the outside ambient heat, light and humidity protected (‘covered’) or non-protected (‘uncovered’) from precipitation for varying lengths of time (Supplementary Table 1). Human skin total RNA was obtained from commercial sources: Stratagene/Agilent Technologies (Santa Clara, CA), Biochain^®^ (Hayward, CA), Zenbio (Research Triangle Park, NC), and Zyagen (San Diego, CA). Human brain total RNA was obtained from a commercial source (Biochain^®^) (run as an internal positive control and not used in any data analysis). Cellular skin samples were collected by swabbing human skin or a touched object surface with a sterile water pre-moistened sterile swab. For all RNA isolations, ½ or a whole 50 μl stain or single cotton swab was used. All samples were stored at-20^o^C until needed, except for the total RNA samples which were stored at-47^o^C.

Suspected bio-particles from male shirt collar samples were collected as previously described [44]. Briefly, WF Gel-Film^®^ x8 retention level (Gel-Pak^®^, Hayward, CA), was cut to a size appropriate for subsequent attachment to a glass microscope slide support (3” x 1” x 1mm, Fisher Scientific, Suwanee, GA). Using sterile tweezers, the back protective covering was removed to expose the adhesive back and the Gel-Film^®^ was placed onto a clean glass microscope slide. The top protective plastic film layer was then removed using re-sterilized tweezers. The Gel-Film^®^ surface was then repeatedly touched to the sample area (direct skin, clothing or object surface) several times to ensure sufficient transfer of biological material. Samples were stained with Trypan Blue (0.4%) (Sigma-Aldrich, St. Louis, MO) for 1 minute, then washed briefly by gentle flooding with sterile ultrapure water with a resistivity of 18.2MΩ at 25^o^C. Samples were then air-dried at room temperature prior to proceeding to sample collection. All samples were stored at room temperature in microscope slide boxes protected from light. Bio-particles were viewed, imaged and collected using a Leica M205C stereomicroscope (Micro Optics of FL, Inc, Davie, FL). Twenty-five, fifty and one hundred bio-particles (i.e. single cells or ‘cellular agglomerates’) were collected. Bio-particles were collected from Gel-Film^®^ surface using 3M™ water-soluble wave solder tape (5414 transparent) on the end of a tungsten needle. The 3M™ water-soluble adhesive was adhered to a clean glass microscope slide using double sided tape and collected on the end of a tungsten needle under the stereomicroscope. The collected bio-particles were then transferred into a sterile 0.2ml PCR flat-cap tube (Phenix Research, Candler, NC)) containing lysis buffer: 100 bio-particle shirt collar sample - 10 μl of lysis buffer solution: 2.1X buffer-blue, 10% *forensic*GEM^TM^ reagent (ZyGEM *forensic*GEM™ tissue kit, VWR, Suwanne, GA), sterile water; 25 and 50 bio-particle shirt collar samples - 5 μl of lysis buffer solution: 1X buffer-silver, 5% *RNA*GEM^TM^ reagent (ZyGEM *RNA*GEM™ tissue kit, VWR), sterile water.

### 2.2 RNA Isolation

Total RNA was extracted from blood, semen, saliva, vaginal secretions, menstrual blood and skin using a manual organic RNA extraction (guanidine isothiocyanate-phenol:chloroform) as previously described [33,45]. Briefly, 500 μl of pre-heated (56^o^C for 10 minutes) denaturing solution (4M guanidine isothiocyanate, 0.02M sodium citrate, 0.5% sarkosyl, 0.1M β-mercaptoethanol) was added to a 1.5mL Safe Lock extraction tube (Eppendorf, Westbury, NY) containing the stain or swab. The samples were incubated at 56^o^C for 30 minutes. The swab or stain pieces were then placed into a DNA IQ^TM^ spin basket (Promega, Madison, WI), re-inserted back into the original extraction tube, and centrifuged at 14,000 rpm (16,000 x g) for 5 minutes. After centrifugation, the basket with swab/stain pieces was discarded. To each extract the following was added: 50 μl 2 M sodium acetate and 600 μl acid phenol:chloroform (5:1), pH 4.5 (Ambion by Life Technologies). The samples were then centrifuged for 20 minutes at 14,000 rpm (16,000 x g). The RNA-containing top aqueous layer was transferred to a new 1.5ml microcentrifuge tube, to which 2 μl of GlycoBlue^TM^ glycogen carrier (Ambion by Life Technologies) and 500 μl of isopropanol were added. RNA was precipitated for 1 hour at-20^o^C. The extracts were then centrifuged for 20 minutes at 14,000 rpm (16,000 x g). The supernatant was removed and the pellet was washed with 900 μl of 75% ethanol/25% DEPC-treated water. Following a centrifugation for 10 minutes at 14,000 rpm (16,000 x g), the supernatant was removed and the pellet dried using vacuum centrifugation for 3 minutes. Twenty microliters of pre-heated (60^o^C for 5 minutes) nuclease free water (Ambion by Life Technologies) was added to each sample followed by an incubation at 60^o^C for 10 minutes. All extracts were DNase treated to remove residual DNA using the Turbo DNA-*free*^TM^ kit (Applied Biosystems (AB) by Life Technologies, Carlsbad, CA) according to the manufacturer’s protocol. With each extraction, a negative control (extraction reagents without sample) was included.

Alternatively, total RNA was extracted from blood, semen, saliva, vaginal secretions, menstrual blood and skin using direct lysis without purification. One hundred microliters of Buffer RLT Plus (QIAGEN, Germantown, MD) with 1 μl β- mercaptoethanol was added to a 1.5mL Safe-Lock extraction tube (Eppendorf, Westbury, NY) containing the stain or swab. The samples were incubated at room temperature for 5 minutes with constant vortexing (20 second intervals). The swab or stain pieces were then placed into a DNA IQ^TM^ spin basket (Promega, Madison, WI), re-inserted back into the original extraction tube, and centrifuged at 14,000 rpm (16,000 x g) for 5 minutes. After centrifugation, the basket with swab/stain pieces was discarded. All samples were stored at-20^o^C until needed.

Total RNA was extracted from bio-particles using the ZyGEM *forensic*GEM^TM^ or *RNA*GEM^TM^ tissue kits (VWR). For the *forensic*GEM^TM^ kit, samples were lysed at 75^o^C for 15 minutes. For the *RNA*GEM^TM^ kit, samples were lysed at 75^o^C for 5 minutes. All samples were stored at-20^o^C until needed.

### 2.3 RNA Quantitation

RNA extracts (manual organic RNA extraction only) were quantitated with Quant-iT^TM^ RiboGreen^®^ RNA Kit (Invitrogen by Life Technologies, Carlsbad, CA) as previously described [33,45]. Fluorescence was determined using a Synergy^TM^ 2 Multi-Mode microplate reader (BioTek^®^ Instruments, Inc., Winooski, VT).

### 2.4 NanoString^®^ Technology

NanoString^®^ standard gene expression chemistry utilizes two ∼50 base probes, the reporter probe and the capture probe, for each mRNA target of interest [43]; when multiplexed, the probe pairs are referred to as a CodeSet. A multiplex CodeSet can be designed to have probe pairs targeting between 20 and 800 mRNAs. Each capture/reporter probe pair within the CodeSet is specifically designed to hybridize to an individual mRNA target. The reporter probe carries the signal and is comprised of a unique molecular fluorescent barcode binding to the 5’ end of the mRNA target. The capture probe binds to the 3’ end of the mRNA target and adheres the capture probe/barcode/target complex to the cartridge surface for data collection (see Figure 1). After overnight hybridization at 65°C in a thermal cycler (typical time of 12-24 hours), the complex is purified on the nCounter Prep Station with excess, unbound probes removed and intact complexes bound, stretched and immobilized on an nCounter Cartridge. Sample cartridges are then placed onto the nCounter Digital Analyzer for counting and data collection of each target complex. The number of times each barcode is counted is proportional to the abundance of that mRNA target in a given sample.

**Figure 1.**
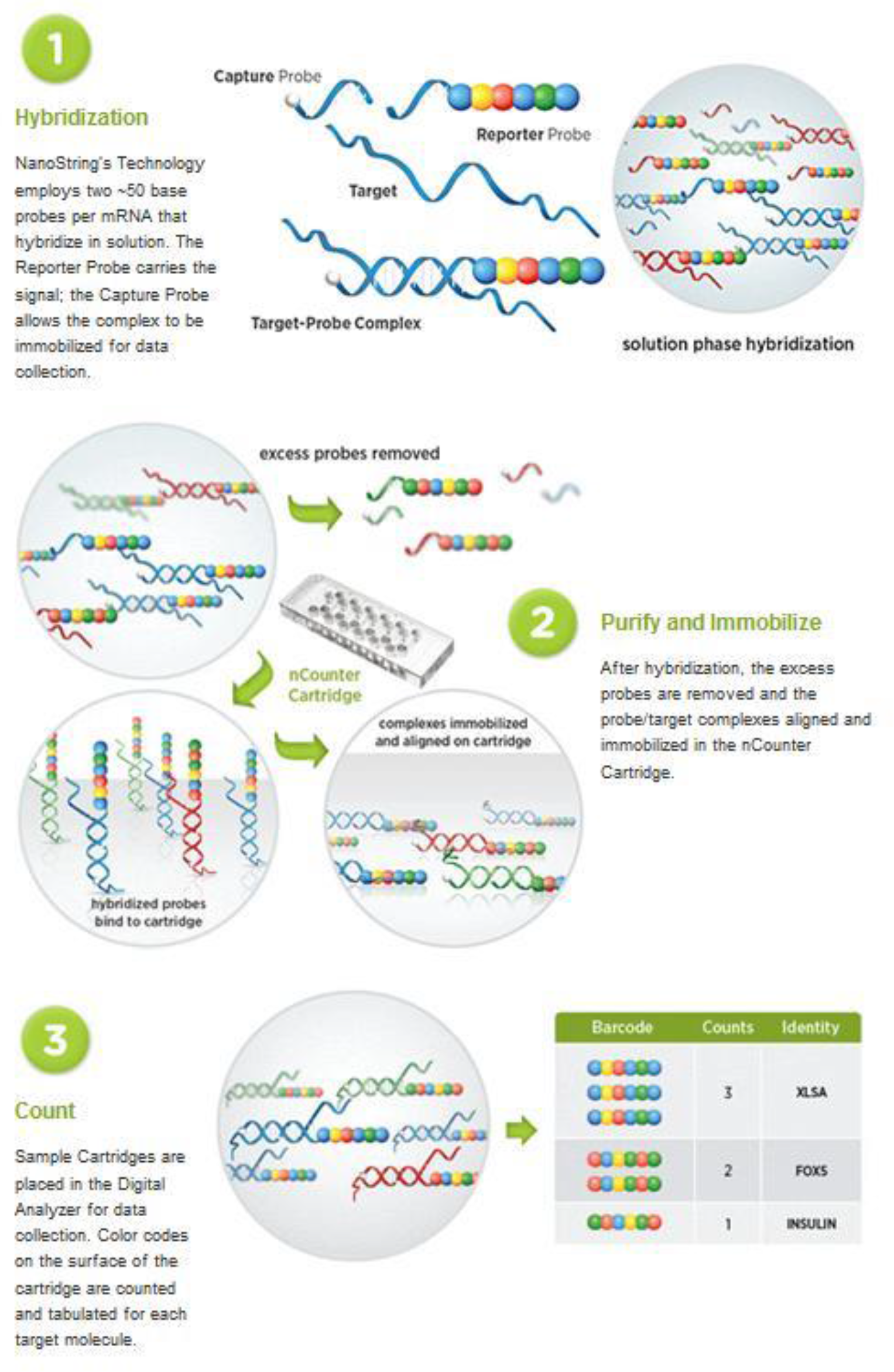
NanoString^®^ digital gene expression technology

In this study, a NanoString^®^ multiplex custom CodeSet was designed and created to target 23 genes known to be differentially expressed in forensically relevant body fluids and tissues. As a reference, 10 ubiquitously expressed housekeeping genes were also included in the CodeSet, giving a 33-plex total. The body fluids and tissues targeted include: venous blood, menstrual blood, semen, saliva, vaginal secretions, and skin. The multiplex CodeSet consisted of 3 venous blood genes (ALAS2, ANK1, HBB) [11,13,34], 2 menstrual blood genes (LEFTY2, MMP10) [15,34,36], 3 saliva genes (HTN3, MUC7, STATH) [3,14,34], 3 semen genes (PRM2, SEMG1, TGM4) [14,34], 5 skin genes (CCL27, IL1F7, KRT9, LCE1C, LCE2D) [33,46], 7 vaginal secretion genes (CYP2A7, CYP2B7P1, DKK4, FUT6, IL19, MYOZ1, NOXO1) [45] and 10 reference (i.e. housekeeping) genes (B2M, COX1, HPRT1, PGK1, PPIH, S15, TCEA1, TFRC, UBC, UBE2D2) (Table 2). The CodeSet also included 6 positive control probes and 8 negative control probes. The 6 positive control probes are designed to assess overall assay performance and to normalize the data, accounting for any assay variability within the system. The 8 negative control probes have no corresponding targets within the sample and assess background noise in the system.

**Table 2.** Body Fluid Specific and Housekeeping Genes in the NanoString^®^ Custom CodeSet

| *Gene* | *Body Fluid Target* |
| --- | --- |
| ALAS2 | Blood |
| ANK1 | Blood |
| HBB | Blood |
| LEFTY2 | Menstrual Blood |
| MMP10 | Menstrual Blood |
| HTN3 | Saliva |
| MUC7 | Saliva |
| STATH | Saliva |
| PRM2 | Semen |
| SEMG1 | Semen |
| TGM4 | Semen |
| CCL27 | skin |
| IL1F7 | skin |
| KRT9 | skin |
| LCE1C | skin |
| LCE2D | skin |
| CYP2A7 | vaginal |
| CYP2B7P1 | vaginal |
| DKK4 | vaginal |
| FUT6 | vaginal |
| IL19 | vaginal |
| MYOZ1 | vaginal |
| NOXO1 | vaginal |
| B2M | Housekeeping Gene |
| COX1 | Housekeeping Gene |
| HPRT1 | Housekeeping Gene |
| PGK1 | Housekeeping Gene |
| PPIH | Housekeeping Gene |
| S15 | Housekeeping Gene |
| TCEA1 | Housekeeping Gene |
| TFRC | Housekeeping Gene |
| UBC | Housekeeping Gene |
| UBE2D2 | Housekeeping Gene |

A total of 96 assays were included this study, involving 89 samples with technical replicates for 7 of the samples. A detailed summary of the 89 samples is provided in Table 1 and includes 14 blood, 17 semen, 17 saliva, 10 vaginal secretions, 10 menstrual blood, and 14 skin samples as well as 5 mixtures and 2 RNA-free controls. For each body fluid both standard and challenging or environmentally compromised samples were evaluated. Full sample descriptions, including number of donors, and the input (ng of total RNA or volume (μl) of extract) used for each of the 96 samples is provided in Supplementary Table 1.

**Table 1.** List of Samples Tested

| **Sample Type** | **N** | **Description** |
| --- | --- | --- |
| ***Blood*** | ***14*** |  |
| Organic Extraction | 7 | Blood stain on cotton cloth (-47°C storage after drying) |
|  | 1 | Environmental (outside (FL) – heat, sunlight, humidity, rain (1 month) |
|  | 1 | Environmental (outside (FL) – heat, sunlight, humidity, covered (3 days) |
| Direct Lysis (RLT) | 5 | Blood stain on cotton cloth (-47°C storage after drying) |
| ***Semen*** | ***17*** |  |
| Organic Extraction | 7 | Dried on cotton swabs (-47°C storage after drying) |
|  | 2 | Environmental (outside (FL) – heat, sunlight, humidity, covered (1 week) |
|  | 3 | Sensitivity: 25ng, 12.5ng, 6.25ng (input achieved by use of 5µl of extract) |
| Direct Lysis (RLT) | 5 | Dried on cotton swabs (-47°C storage after drying) |
| ***Saliva*** | ***17*** |  |
| Organic Extraction | 7 | Dried buccal sample on cotton swabs (-47°C storage after drying) |
|  | 1 | Environmental (outside (FL) – heat, sunlight, humidity, rain (1 week) |
|  | 1 | Environmental (outside (FL) – heat, sunlight, humidity, covered (1 month) |
|  | 3 | Sensitivity: 25ng, 12.5ng, 6.25ng (input achieved by use of 5µl of extract) |
| Direct Lysis (RLT) | 5 | Dried buccal sample on cotton swabs (-47°C storage after drying) |
| ***Vaginal Secretions*** | ***10*** |  |
| Organic Extraction | 6 | Dried sample on cotton swabs (-47°C storage after drying) |
|  | 1 | Environmental (outside (FL) – heat, sunlight, humidity, rain (3 days) |
| Direct Lysis (RLT) | 3 | Dried sample on cotton swabs (-47°C storage after drying) |
| ***Menstrual Blood*** | ***10*** |  |
| Organic Extraction | 7 | Dried sample on cotton swabs (-47°C storage after drying) |
| Direct Lysis (RLT) | 3 | Dried sample on cotton swabs (-47°C storage after drying) |
| ***Skin*** | ***14*** |  |
| Organic Extraction | 1 | Swab of surface skin (male hand); swab moistened with sterile water |
|  | 1 | Swab of coffee cup surface; swab moistened with sterile water |
|  | 1 | Swab of computer mouse; swab moistened with sterile water |
| Direct Lysis (RLT) | 1 | Swab of surface skin (male hand); swab moistened with sterile water |
|  | 1 | Swab of coffee cup surface; swab moistened with sterile water |
|  | 1 | Swab of computer mouse; swab moistened with sterile water |
| Direct Lysis (RNAGEM) | 1 | 25 bio-particles (clumps); shirt collar (male) |
|  | 1 | 50 bio-particles (clumps); shirt collar (male) |
| Direct Lysis (*forensic*GEM) | 1 | 100 bio-particles (55 clumps/45 singles); shirt collar (male) |
| None | 5 | Skin total RNA (commercial source) |
| ***Mixtures*** | ***5*** |  |
| Organic Extraction | 2 | Vaginal/semen (1/2 swab of each donor extracted in same tube) |
|  | 2 | Blood/saliva (1/2 stain/swab of each donor extracted in same tube) |
|  | 1 | Semen/saliva/vaginal (1/2 swab of each donor extracted in same tube) |
| ***Controls*** | ***3*** |  |
| Organic Extraction | 2 | Clean sterile swab (negative control) |
| None | 1 | Brain total RNA* (commercial source) |

Stain = 50 µl stain; Swab – saturated body fluid swab (sterile cotton)
Environmental samples (blood, semen, saliva) – on cotton cloth
Total RNA – commercial sources (see methods)
\* run as an internal positive control and not used in any data analysis

Hybridization assays were performed according to the standard NanoString^®^ gene expression assay protocol, as follows: Each individual assay consisted of 10µL Reporter Probe, 10µL Hybridization Buffer, 5µL Capture Probe and the specified RNA sample input (in most cases, 50ng of total RNA or 5µL crude lysate) for a total reaction volume of 30µL. Assays were placed into a thermal cycler at 65°C with a 70°C lid, and allowed to hybridize overnight for approximately 16 hours. Following this, assays were placed onto the nCounter Prep Station using the high-sensitivity protocol for purification and immobilization of the hybridized targets on the imaging cartridge. The cartridges were then scanned on the nCounter Digital Analyzer for counting of the hybridized targets, and data files were exported for analysis.

### 2.5 Statistical Methods

#### 2.5.1 Overview of method

Our approach to the problem is motivated by three properties of *bona fide* casework samples: they often (i) comprise mixtures of two or more fluids, (ii) are limited in quantity and (iii) could be either partially or highly degraded. Our basic approach is as follows: First, we model the probability distribution of gene expression in body fluid samples. Next, we use this model to calculate the Maximum Likelihood Estimate (MLE) for the levels of each body fluid in a sample and to calculate the log-likelihood of a sample’s profile given the estimated levels of each fluid. We then construct a likelihood ratio comparing the likelihood of a given sample’s profile with and without the presence of a given fluid. If a sample’s profile is far more likely when we include a specific fluid in the model, then we conclude the fluid is present in the sample.

#### 2.5.1 Modeling gene expression in body fluids

Gene expression is best modeled on the log (multiplicative) scale: a doubling of a gene’s expression level is generally considered a change comparable in magnitude to a halving of its expression level, and a gene increasing from 200 to 400 mRNA transcripts is as meaningful a difference in gene expression as a gene increasing from 2000 to 4000 counts. However, the mathematics of mixtures is additive: if a sample is half blood and half saliva, a gene’s cumulative expression level will result from the summation of its expression levels in each tissue. We therefore model the contributions of each fluid to a mixture on the linear scale, but we measure discrepancies between observed and predicted expression on the log scale.

We develop the algorithm as follows: As a conceptual starting point, we first describe a model for gene expression in a sample from a single fluid. We then extend this model to mixtures of fluids. From there we describe calculation of maximum likelihood estimates (MLEs) of fluid quantities in a sample, and we describe the use of likelihood ratios to test for the presence of a fluid in a sample.

#### 2.5.2 Model for gene expression in a sample from a single body fluid

On average, each gene represents a given proportion of total gene expression in each fluid. For example, in the average blood sample we might expect 15% of total RNA to be HBB, 1% to be ALAS2, etc. Call these expected proportions X_HBB_, X_ALAS2_, etc. Then in a given blood sample, the vector of expected gene expression is β(X_HBB_, X_ALAS2_,…)^T^, where β is the total amount of RNA in the sample.

Due to both biological and technical noise, actual expression will vary around its expectation. Per the multiplicative nature of gene expression, we model this variability as arising from a log-normal distribution, and we assume that each gene is equally variable. A single gene’s expression in a sample can then be modeled:

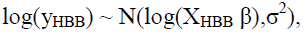

where y_HBB_ is the expression of HBB in the sample, and σ^2^ is the variance (on the log scale) of HBB’s expression around its expectation.

#### 2.5.3 Model for gene expression in mixtures of body fluids

The model for mixtures follows naturally from the model for single-fluid samples. First, let us define notation. We represent matrices with bold, uppercase letters, vectors with bold, lowercase letters, and scalars with lowercase letters. We index samples i∊ (1,…, n), genes j ∊ (1,…, p), and tissues k ∊ (1,…, K). Call the gene expression profile for a given sample **y**_i_ = (y_i1_,…, y_ip_)^T^, where y_ij_ is the expression of gene j in sample i. Call β_ik_ the amount of fluid k in sample i, and call **β**_i_ = (β_i1_,…, β_iK_) the vector of the amounts of all the fluids in sample i. Finally, define the matrix **X** to represent the expected proportion of each gene in each fluid type, with x_jk_, the element in the j^th^ row and the k^th^ column of **X**, representing the expected proportion of gene j in samples from fluid k.

Assuming the number of mRNA molecules in mixtures of fluids will be a sum of the number of mRNA molecules in each component of the mixture, we can write the expected counts of gene j in sample i:

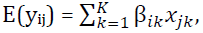

and the expression for the sample’s entire expected gene expression vector is simply

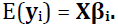

Again assuming the variability of gene expression occurs on the log scale, we model gene expression in a sample as:

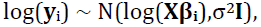

where **I** is the identity matrix and σ^2^ is the common variance (on the log scale) of all genes. (Note that if E(**y**i) = **Xβ**_i_, then E(log(**y**_i_)) ≠ log(**Xβ**_i_). However, under the values considered in this application, E(log(**y**_i_)) very closely approximates log(**Xβi**).) As we lack the data to fully estimate the genes’ covariance matrix, we approximate it with σ^2^**I**.

Before we can apply the above model for gene expression in body fluids, we must estimate two parameters: **X**, the matrix of expected proportions of gene expression, and σ^2^, the variance of gene expression. Estimation of the **X** matrix is described in Section 3.2. We estimated σ^2^, the variance on the log scale common to all genes, as the average variance of each gene in each tissue or fluid.

#### 2.5.4 Maximum likelihood estimation of the amounts of each tissue or fluid in a sample

Under the assumptions that log gene expression is normally distributed around the log of its expectation and that each gene is equally variable, the MLE for **β_i_** can be calculated as follows:

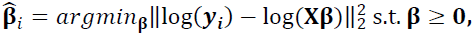

i.e. 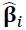 minimizes the sum of squared errors on the log scale between the observed gene expression **y**i and the predicted gene expression **Xβ**, subject to the constraint that all the elements of **β** are non-negative (a sample cannot have negative amounts of a fluid). As it is doubtful that a closed-form solution to this expression exists, we use numerical methods to optimize it [47]. The expression is not convex in **β**; however, we find its estimates to be reasonably robust to differing initial conditions, returning similar estimates with very similar log-likelihoods.

To prevent the algorithm from overexerting itself trying to fit gene expression values in the background of the assay, we found it necessary to add one layer of complexity to the model: in addition to fitting β terms for each fluid, we added a β for background, with a corresponding column in the **X** matrix with equal weights on all genes. We further constrained this background β term to contribute no more than 15 counts to each gene. For the same reason, we truncated all gene expression values at 5 counts, a reasonable estimate of the average background counts.

#### 2.5.5 Using likelihood ratios to test the presence of tissues or fluids

In any given sample **y**_i_, our goal is to determine which tissues or fluids are present. That is, we want to test whether each element of **β_i_** equals 0. A reasonable approach to this problem is to calculate the likelihood of the data under the MLE 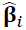 and under a constrained MLE 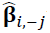 with the β_ij_ term corresponding to the tissue or fluid in question forced to 0. The likelihood ratio under the full and constrained MLEs will summarize the evidence for the presence of the tissue or fluid in question.

Calculation of a log likelihood for the data given a MLE is straightforward. Under our model, log gene expression is normally distributed around the log of the predicted gene expression. Then up to a constant, the log-likelihood of **y**_i_ given 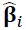 is:

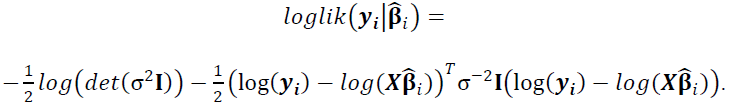

To test whether fluid j is present in sample i, we evaluate the above expression using **y**_i_ and 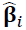 and again using **y**_i_ and the constrained MLE 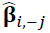, and we calculate a likelihood ratio.

## 3. Results

### 3.1 Selection of mRNA biomarkers

We designed a ‘CodeSet’ to probe 23 body fluid/tissue specific genes and 10 housekeeping genes (Table 2), which is well within the 800 target technological capability of the system. To take advantage of the high multiplex capability of the system, we deliberately included biomarkers that have been demonstrated to be highly specific to a particular body fluid (e.g. PRM2 and SEMG1 for semen) as well as some that have shown a lesser degree of tissue specificity (e.g. MYOZ1 for vaginal secretions and MUC7 for saliva).

### 3.2 Estimating expected body fluid profiles

Our algorithm requires accurate estimates of each fluid’s average gene expression profile; below, we describe the results of this analysis.

Our dataset included samples of highly varying RNA concentration, and genes in the lower-concentration samples frequently dropped into the background noise of the assay. To ensure accurate estimates of each body fluid’s average gene expression profile, we only used samples with high expression levels of housekeeping genes. As a set of ‘training samples’ we took the four highest-expressing samples from each fluid type with the exception of saliva, where a lack of high-expressing samples limited us to three training samples. Supplemental Figure 1 shows the overall housekeeping gene expression levels in the training samples and the remaining samples.

Per our model described in Section 2.5.3, we are interested in the relative expression levels of the genes within each body fluid; that is, in the proportion of total signature gene expression expected from each gene in a given body fluid. (This is in contrast to most gene expression-based classifiers, which are more interested in each gene’s absolute expression level. Since it is unrealistic to expect a housekeeping gene to be invariant across body fluid types, normalizing our data to attain “absolute” expression levels is impossible.) Therefore, we globally normalized each sample, rescaling them so the sum of all expression values was 1 and so that each gene’s expression value was its proportion of the total signature gene expression. We then estimated each gene’s expected proportion of expression in each fluid with its mean normalized expression value within each fluid.

The five body fluids and skin demonstrated highly distinct gene expression profiles, and although the signature genes varied between samples of the same fluid, their differences between fluids were much greater.

Figure 2 shows the expected proportion of total expression for each gene in each fluid. Supplemental Figure 2 shows the consistency of these profiles in the training data, and Supplemental Figure 3 organizes the information in Figure 2 by gene rather than by fluid. In all fluids the average expression profile exhibits elevated expression of the fluid’s putative characteristic genes, although this trend was distinctly weaker in saliva samples.

**Figure 2.**
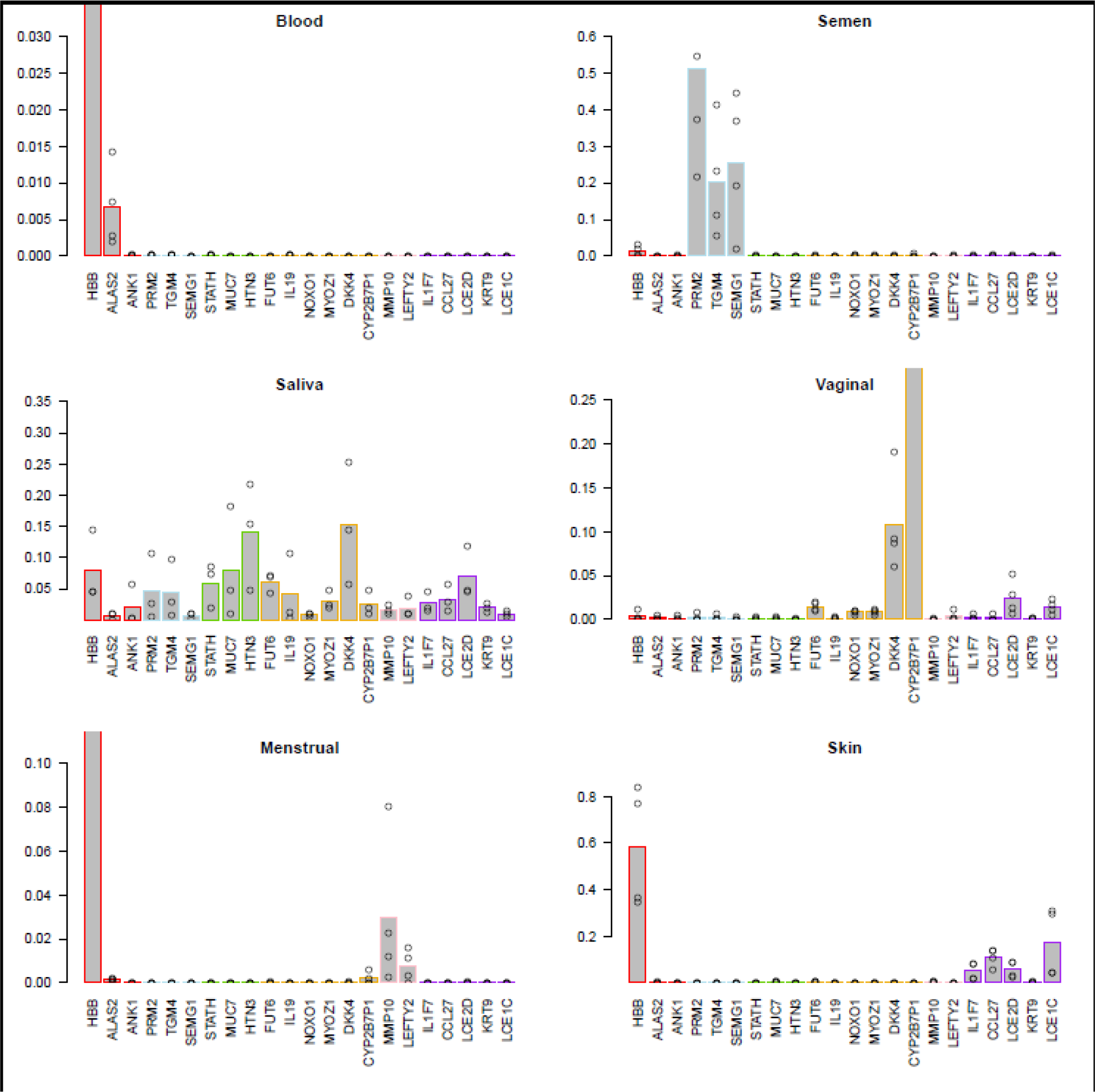
Average proportion of total expression for each gene in each fluid. The vertical axis shows the relative proportion of total gene expression attributable to each gene (on the log scale). For each fluid, each point shows a gene’s relative expression in a single training sample, and each bar shows the average of the gene’s relative expression across the fluid’s training samples. Bar color indicates genes’ putative tissues.

**Figure 3.**
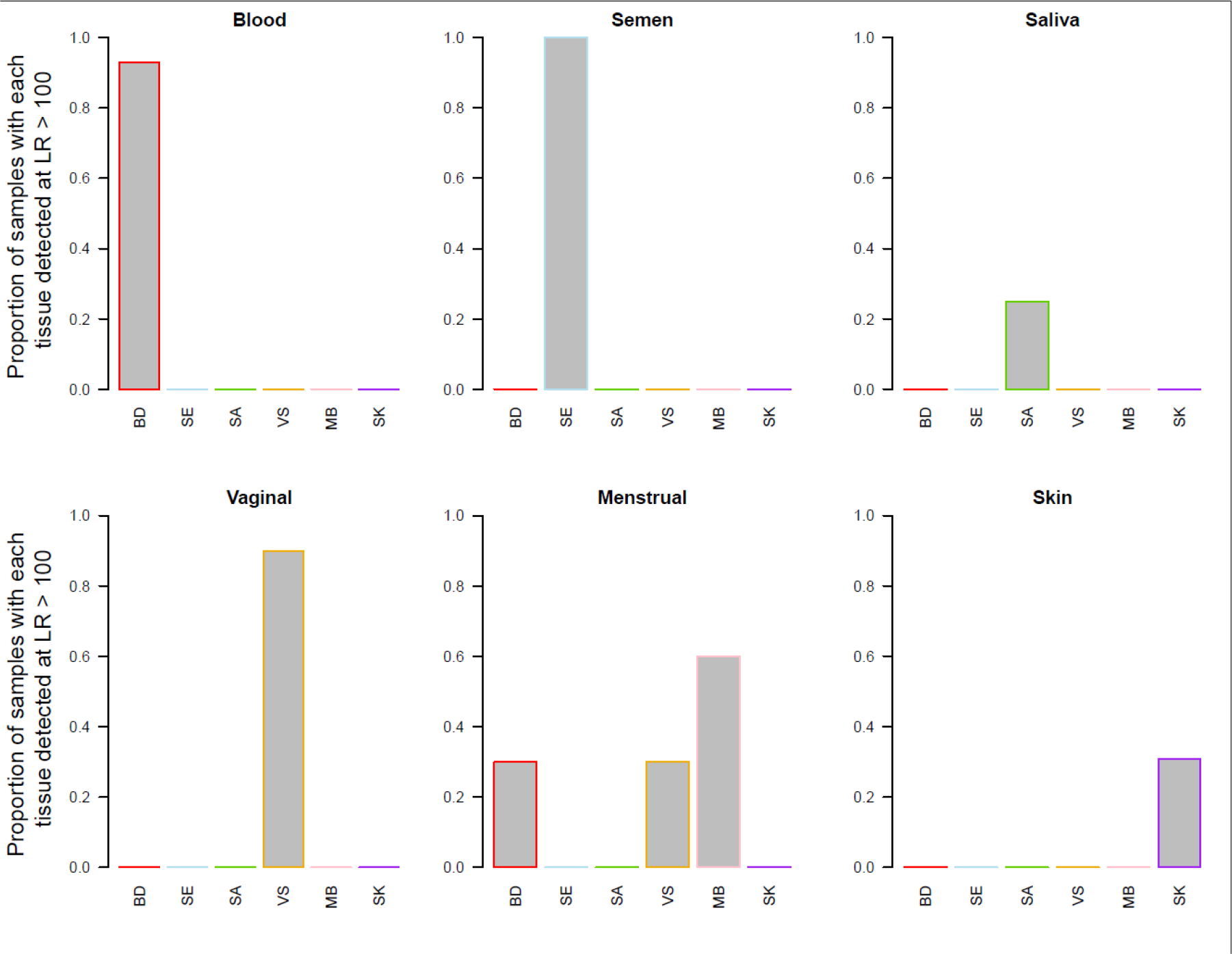
Performance of the algorithm on all single-source samples. Bars display the rate at which each fluid is called detected in each sample type. Fluids are called detected if their likelihood ratio exceeds 100.

HBB expression dominated the blood profiles, far exceeding the other blood markers ALAS2 and ANK1, although ALAS2 levels in blood greatly exceeded those of other genes. The putative blood marker ANK1 was not enriched in blood samples, surprisingly appearing most prominently in saliva samples instead. Expression in semen samples came almost entirely from the semen-specific genes PRM2, TGM4 and SEMG1, although other genes, particularly HBB, were detectable. Saliva samples had the most diffuse profile, with the saliva-specific genes STATH, MUC7 and HTN3 contributing only 28% of total measured expression. Vaginal secretion samples had highly elevated levels of the vaginal markers DKK4, CYP2B7P1 and to a lesser extent FUT6. Menstrual blood samples alone showed elevated expression of their characteristic genes MMP10 and LEFTY2. Unsurprisingly, menstrual blood samples also contained blood (HBB, ALAS2) and vaginal secretion (CYP2B7P1) biomarkers. Skin samples showed elevated expression of the skin genes LCE1C, IL1F7 and CCL27, although these genes were also slightly elevated in vaginal secretions and menstrual blood. HBB was the most prevalent gene in the commercial skin preparation, probably due to the inevitable presence of contaminating endothelial tissue in such preparations.

Most genes were present at a non-negligible proportion of total expression in the saliva samples. This phenomenon results from this study’s lack of a good saliva marker. If a gene highly expressed in saliva were measured, the relative expression of the other fluids’ characteristic genes in saliva would shrink dramatically.

### 3.3 Using gene expression to predict the body fluid composition of samples

Our algorithm for body fluid detection is described in detail in the Methods section. Below, we summarize the performance of the method in predicting the body fluid composition of every sample in our study. Crucially for forensic applications, our test appears to have extremely high specificity; in fact, it returned zero false positives in this study’s 89 samples.

We used a likelihood ratio cutoff of 100 to declare whether a body fluid was detected in a given sample, and found that 53/80 single-fluid, non-duplicate samples (66%) gave positive results. It is worth noting that our collection of samples was not necessarily representative of the real world population of forensic samples, as in many cases we intentionally chose degraded and miniscule samples to push the limits of the assay. Figure 3 shows the rate at which each body fluid was declared ‘detected’ in each actual fluid using an LR of 100. Supplemental Figures 4 and 5 indicate the performance of the algorithm in the training samples (abundant RNA) and in the remaining samples (low RNA quantity) respectively. The algorithm was successful in identifying the correct body fluid as long as the sample was abundant enough; in low input samples it detected blood, semen and vaginal secretions reliably while struggling to detect saliva, menstrual blood and skin. Across all samples, detection of blood, semen and vaginal secretions was nearly perfect. Menstrual blood was successfully detected 60% of the time. Blood and vaginal secretions were frequently detected in menstrual blood, though these cannot be considered false positives. Rather, it appears menstrual blood is best modeled as a variable mixture of blood, menstrual blood, and vaginal secretions. Saliva was successfully detected in only 25% of samples, likely due to fact that the characteristic saliva genes were not as informative as other fluids’ characteristic signature genes and/or to the very low level of total RNA in most of the saliva samples. Skin also proved difficult to detect (31% success rate); however, the need to identify skin will probably be limited to specialized forensic cases. It is much more important to ensure that skin samples are not misclassified as other tissues.

**Figure 4.**
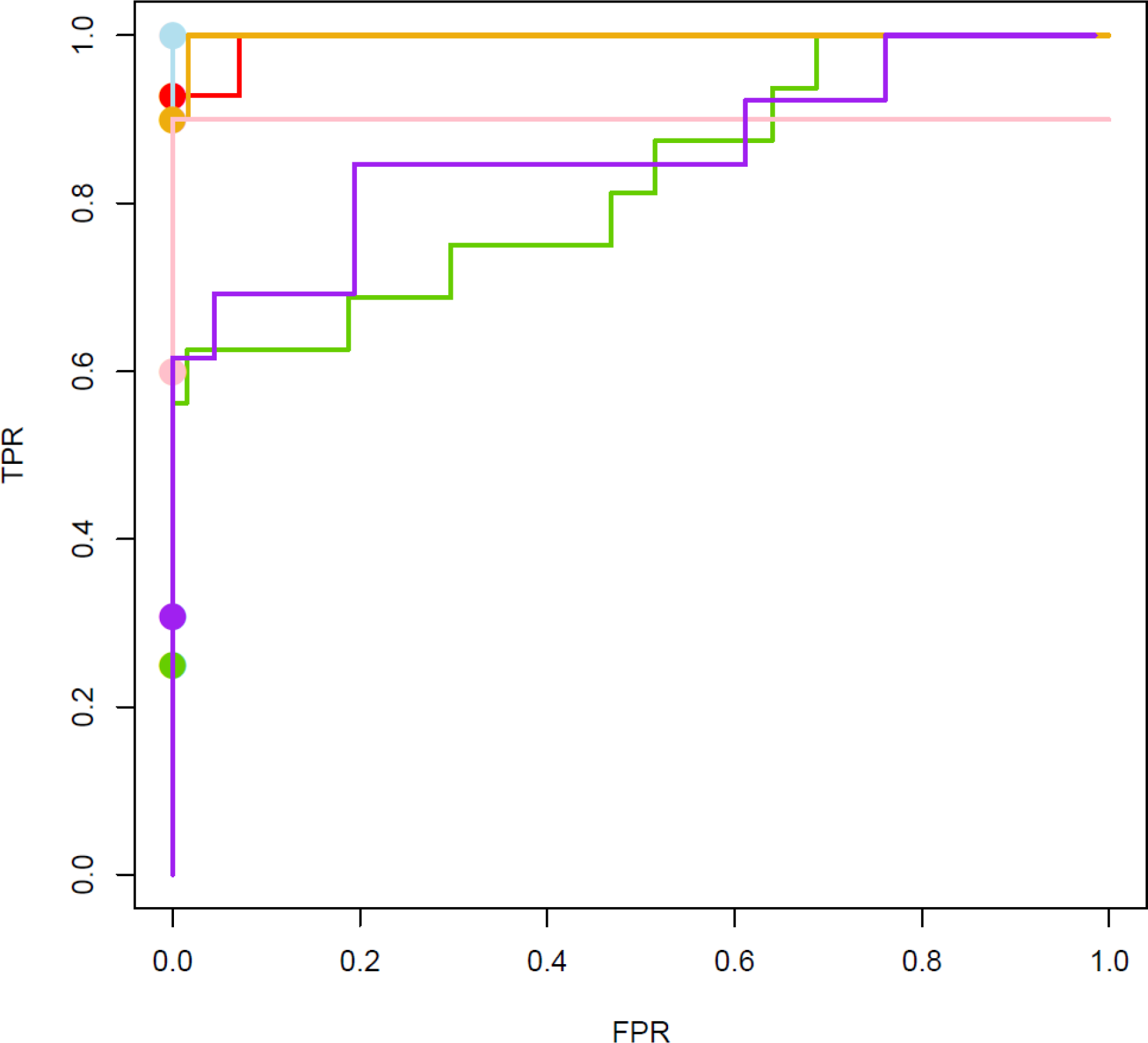
ROC curves showing the algorithm’s True Positive Rate (TPR) and False Positive Rate (FPR) for each tissue.

Points indicate the performance achieved using a LR cutoff of 100. Relaxing this LR cutoff for detection of menstrual blood, saliva and skin could greatly increase the TPR without increasing the FPR. Line color indicates body fluid: blood – red, semen – blue, saliva – green, vaginal – orange, menstrual blood – pink, skin – purple.

**Figure 5.**
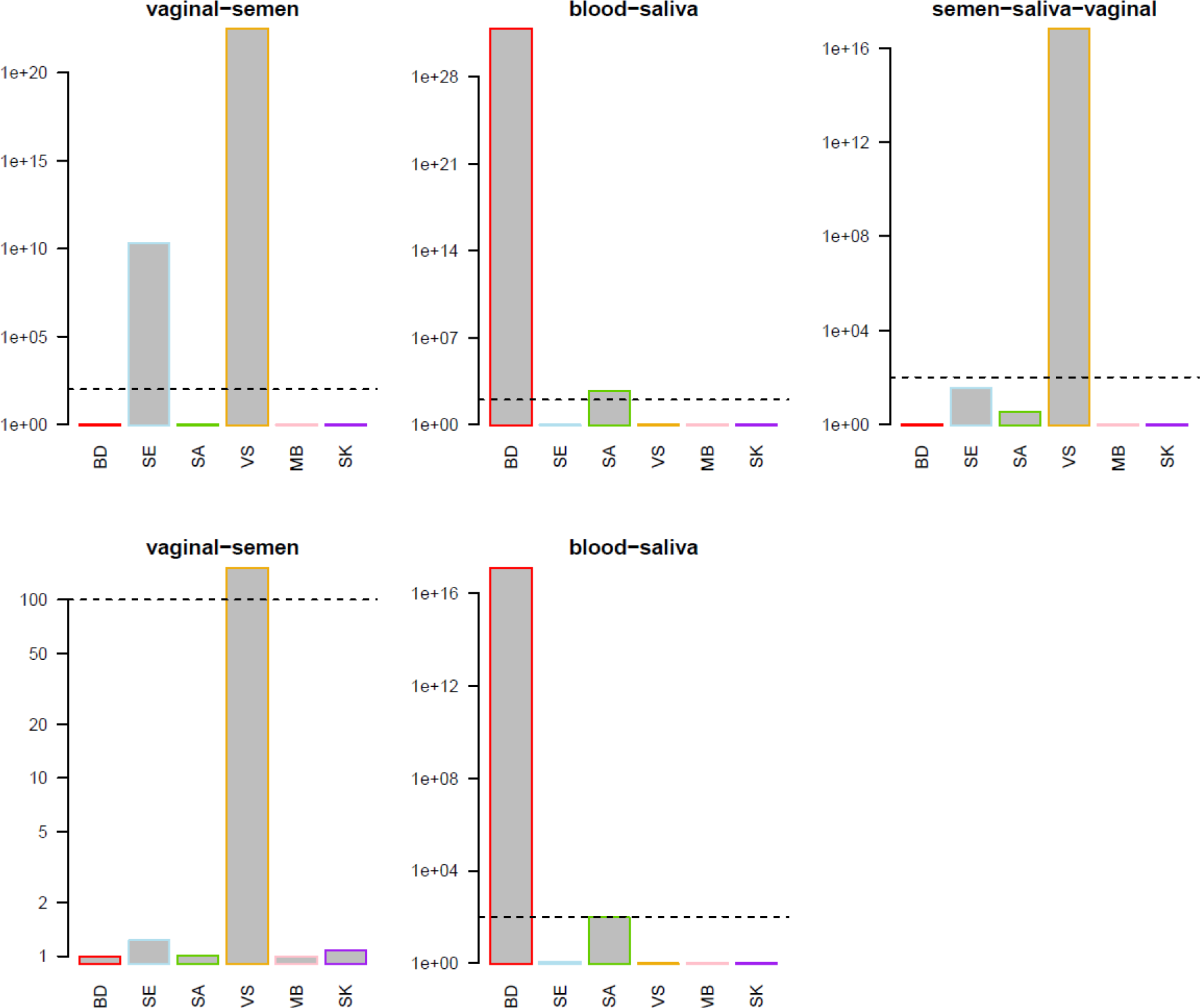
Performance of the algorithm in five mixture samples. For each of five mixture samples, a bar plot shows the likelihood ratios for the presence of each fluid type. The dotted line indicates a LR of 100.

The choice of a LR >100 cutoff for detecting fluids is arbitrary, and our algorithm could achieve better performance with a less strict cutoff. Figure 4 shows ROC curves for the True Positive Rate (TPR) and False Positive Rate (FPR) for detection of each fluid type in our data. As the LR threshold relaxes our test returns more of both false positives and false negatives. For the tissues with the worst performance in our data – menstrual blood, saliva and skin – the ROC curves reveal that a relaxation of the LR thresholds in some tissues would result in large increases in TPR without any increase in FPR.

### 3.4 Body fluid mixtures

As a preliminary indication of the ability of the method to discern admixtures of body fluids, five mixtures were prepared by combining ½ of a 50 μl stain or single cotton swab from each body fluid. The mixtures comprised four binary (2 × vaginal secretions/semen, 2 × blood/saliva) and one ternary mixture (semen/saliva/vaginal secretions). The blood/saliva and vaginal secretions/semen were biological, as opposed to technical, replicates since the donors were different. Using an LR of 100 as a decision threshold, two of the five mixtures were called perfectly, namely one of the vaginal secretions/semen and one of the blood/saliva samples (Figure 5). One of the component fluids was identified in each of the three ‘false negative’ mixtures: vaginal secretions (vaginal secretions/semen and semen/saliva/vaginal secretions) and saliva (blood/saliva). In the latter ternary mix the semen and saliva components were detected but with LRs of <100 (36.9 and 3.4 respectively). In the second blood-saliva sample, the LR for saliva was 95, falling just short of our strict bar for detection. In all but one of the mixture samples, the component fluids are evident from their likelihood ratio profiles: using an LR cutoff of 5, four of the five mixtures were called perfectly. Significantly, no false positives were observed even under the very generous LR cutoff of 5.

### 3.5 Development of a routine-use 5 minute RNA direct lysis method

To facilitate routine analysis, we tested a simple 5 minute room temperature cellular lysis protocol as an alternative to standard RNA isolation for forensic sample processing using the NanoString^®^ procedure (See Methods Section). The method is based upon the RLT buffer from QIAGEN which contains a high concentration of guanidine thiocyanate as well as a proprietary mix of detergents. β-mercaptoethanol (1% v/v) is also added before use to inactivate RNAses in the lysate. The NanoString assay involves direct hybridization to the RNA with no enzymatic steps, and neither the presence of the denaturing buffer nor the cellular debris in the lysate have a significant impact on the assay results.

We compared the reproducibility of the assay between standard RNA isolation/purification and direct lysis protocols from the same source material. Fourteen of the samples in our study were compared in this manner. Supplemental Figure 6 shows scatterplots comparing log expression values for each of these same source samples between the two protocols. In general we saw excellent concordance between the two protocols for all genes with a moderate to high degree of expression. The correlation between the protocols breaks down for very lowly-expressed genes, reflecting the greater noise in the assay when measuring vanishing target. The most dramatic differences between replicates (for example in the samples MB-2 and BD-5) are attributable to expected variance in RNA input amounts between lysate and purified RNA since lysate concentration is not reliably measureable by current methods. The concordance observed between lysis and purified protocols suggest that the simpler, 5 minute lysis protocol would be an efficient option for routine forensic casework workflow.

## 4. Discussion

The results of this preliminary proof of principle study indicate that it is feasible to identify the common forensically relevant body fluids by multiplex solution hybridization of barcode probes to specific mRNA targets using a simple five minute direct lysis protocol. This simplified protocol with minimal hands-on requirement should facilitate routine use of mRNA profiling in casework laboratories. We first describe a model for gene expression in a sample from a single body fluid and then extend that model to mixtures of body fluids. From there we describe calculation of maximum likelihood estimates (MLEs) of body fluid quantities in a sample, and we describe the use of likelihood ratios (LR) to test for the presence of each body fluid in a sample. In contrast to most gene expression-based classifiers, we do not train a machine learning algorithm to optimize our ability to call samples correctly; rather, we define a biologically reasonable model of gene expression in body fluid samples and we use that model to evaluate the strength of evidence a sample provides for the presence of a particular fluid. This founding of our algorithm in sound statistical principles allows the calculation of log-likelihoods for detection of each fluid type, making the algorithm’s results defensible in courtroom settings.

A further benefit of this principled approach is that it allows us to evaluate our algorithm on all samples, including those used in training: as our algorithm is based on an *a priori* model of gene expression in body fluid mixtures, and as we estimated its parameters without regard to model performance, the algorithm can only minimally overfit the training data. Our algorithm’s performance in the training samples may therefore slightly overestimate its performance in future samples, while its performance in the other, low-RNA samples will considerably underestimate future performance in high-quality samples. Although we initially used an LR of 100 as the decision threshold for all body fluid types, we subsequently demonstrated that it may be possible to use a less restrictive threshold to improve the positive call rate without generating false positives. Alternative approaches using body fluid-specific thresholds should be investigated.

While the prototype biomarker ‘CodeSet’ performed remarkably well in the work described herein, further optimization of the biomarkers will be required before the method can be used in casework. The HBB blood biomarker is approximately 1000-fold more highly expressed than ALAS2, the second-most prevalent blood marker in our data. This means that HBB’s limit of detection (LOD) is so low that the possibility of false positives with non-blood body fluids increases due to possible low level contamination with vascular tissue products. This potentially confounding issue can be addressed by attenuating the HBB signal with the addition of precisely defined quantities of specifically designed unlabeled oligonucleotides complementary to the HBB RNA prior to hybridization with the full CodeSet. These competitively inhibit the hybridization reaction with the labeled probes.

In contrast to the need to attenuate one of the blood biomarkers, the signal for the saliva biomarkers needs to be enhanced. The most specific and highly expressed saliva biomarker is HTN3. Signal intensification could be accomplished by designing multiple probes that bind along a single HTN3 mRNA. In addition the current probes could be designed to hybridize to both HTN3 and HTN1, the latter of which is also saliva specific. Alternative novel biomarkers identified by RNA-Seq studies could also be employed if the HTN3 intensification strategies fall short of expectations.

Some of the selected biomarkers did not perform as expected. For example, the ANK1 blood biomarker did not demonstrate blood specificity in the NanoString^®^ assay with this sample set since the expression level was low in all tissues. Re-design of some probe sequences may be worthwhile, but it is likely that assay performance would be most significantly improved by the incorporation of additional body fluid specific biomarkers (e.g. commensal bacteria from the vagina, such as *Lactobacillus sp.*). Future iterations of the CodeSet will evaluate the performance of additional genes.

As a preliminary indication of the ability of the method to discern admixtures of body fluids, one ternary and four binary mixtures were prepared. The true fluid composition in four of the five mixtures was clear from their likelihood ratio profiles, and at least one fluid was correctly detected in all mixtures. Although these results were encouraging, a thorough investigation of the performance of a more optimized NanoString^®^ assay with a variety of different mixtures will be necessary.

There needs to be a note of caution with respect to the skin assay results. The chosen skin biomarkers were selected using total skin RNA from commercial sources due to the difficulties in isolating sufficient quantities of total RNA from touch samples to perform the hundreds of assays required for the biomarker screening and confirmation process. It is likely that the highly purified commercial skin samples will contain mRNAs that originate from multiple layers of skin including both dermal and epidermal tissue as well as contaminating endothelial tissue and its contents (i.e. blood), and it is likely that *bona fide* touch samples, which presumably mainly consist of cortical cells from the epidermis, will possess a different gene expression profile than that obtained from the commercial product. Some of the putative skin biomarkers were found in some of the other tissues, especially saliva (CCL27, LCE2D, IL1F7, KRT9), a finding perhaps due to common biomarker functions in skin and the alimentary tract or to the presence of skin cells in saliva. The highly expressed blood marker HBB was present in the commercial skin RNA preparations at comparable or higher levels than the highly expressed skin biomarker LCE1C, confirming the presence of contaminating endothelial tissue. In light of the extremely low abundance of tissue in most touch skin samples, it remains to be seen the degree to which skin biomarkers prove generally useful in forensic investigations. We suspect the inclusion of skin-specific genes will at a minimum help forensic assays avoid misclassification of skin samples as other tissues.

Housekeeping genes are typically added to gene expression assays to indicate that RNA of sufficient quality and quantity for analysis is present, and for normalization purposes [6,15,38]. Due to non-uniform expression of housekeeping genes their value as normalizers is questionable [48,49]. Here we show that the developed algorithm does not require normalization with housekeeping genes. However their presence indicates the recovery of suitable RNA for analysis and therefore still has a certain utility in the assay.

## Acknowledgements

The authors would like to acknowledge all of the anonymous donors who provided samples for this study. Support for portions of this work was provided by the State of Florida through the National Center for Forensic Science at the University of Central Florida. The opinions, findings and conclusions or recommendations expressed in this publication are those of the authors and do not necessarily reflect those of the State of Florida.

---

**Supplementary Table 1.**
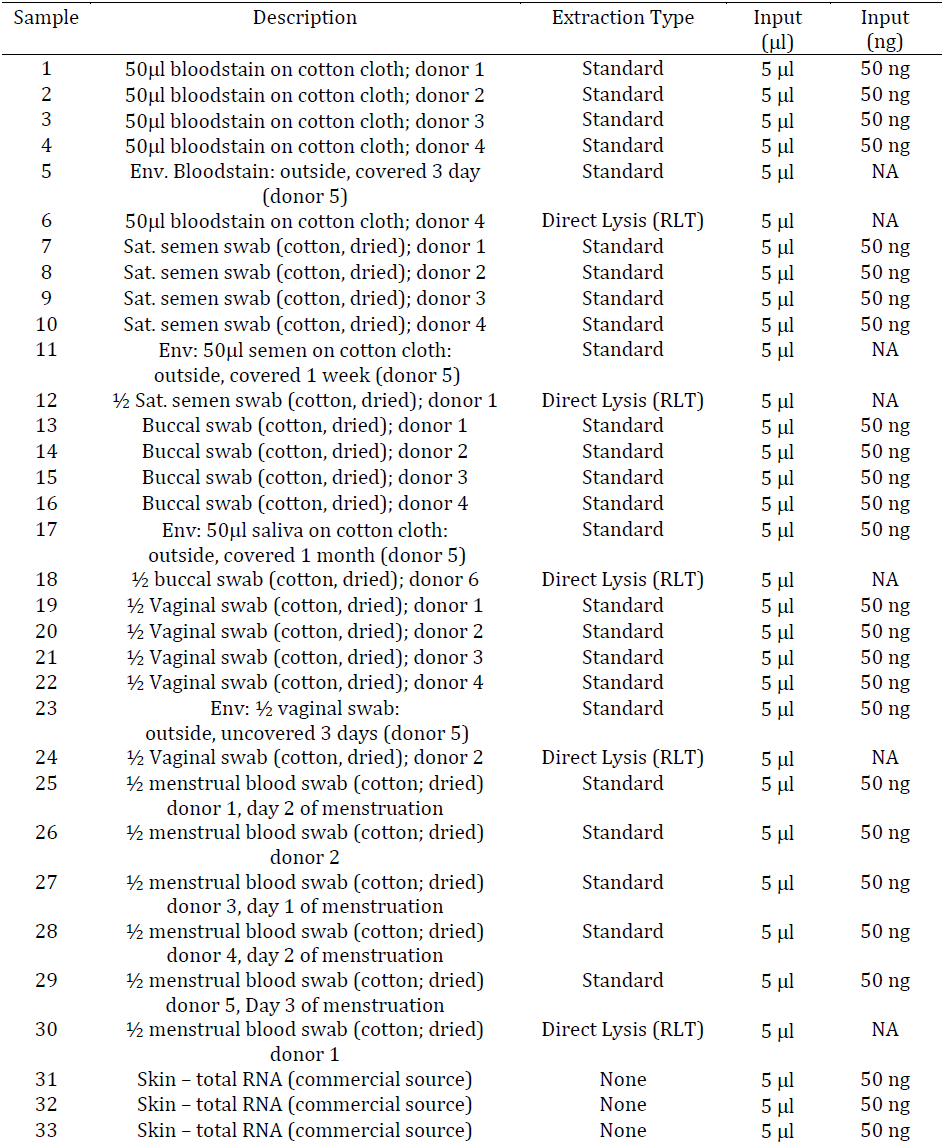
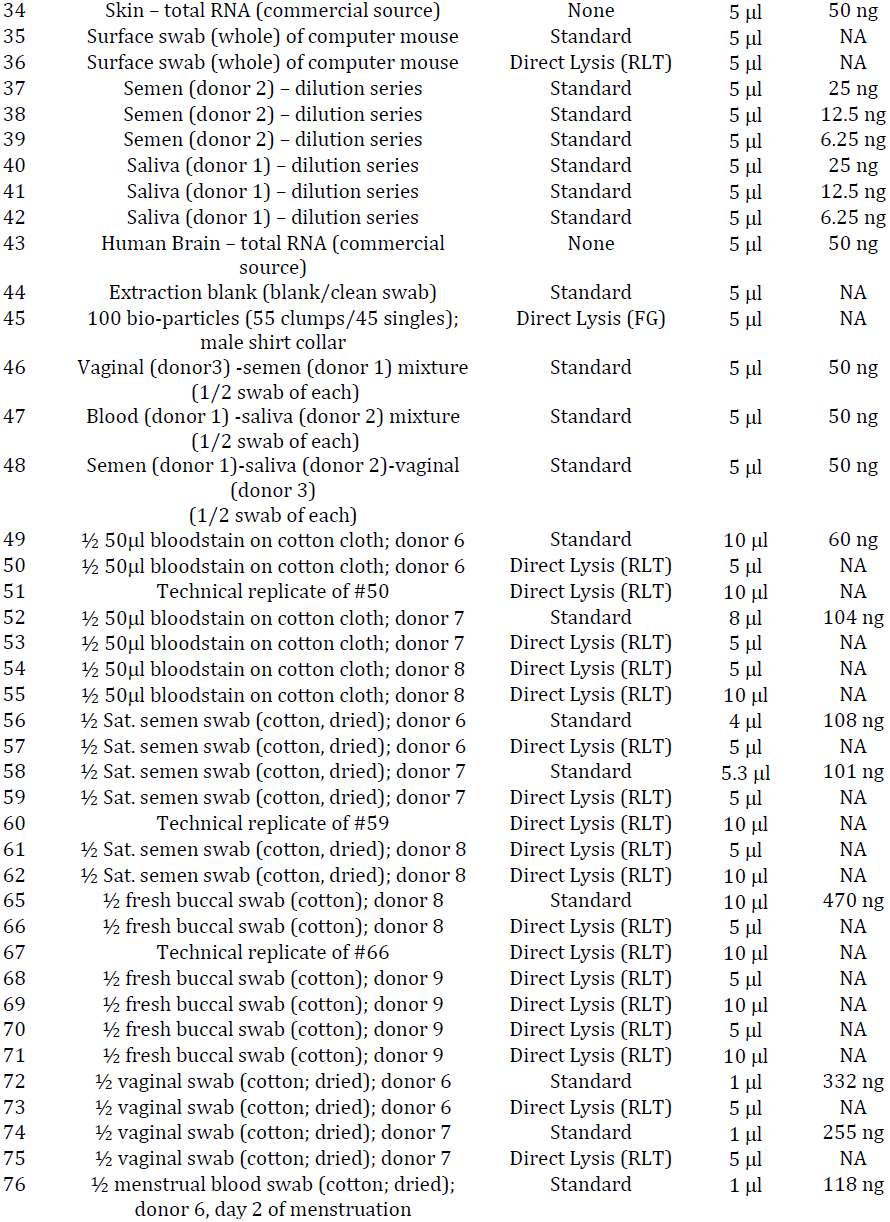
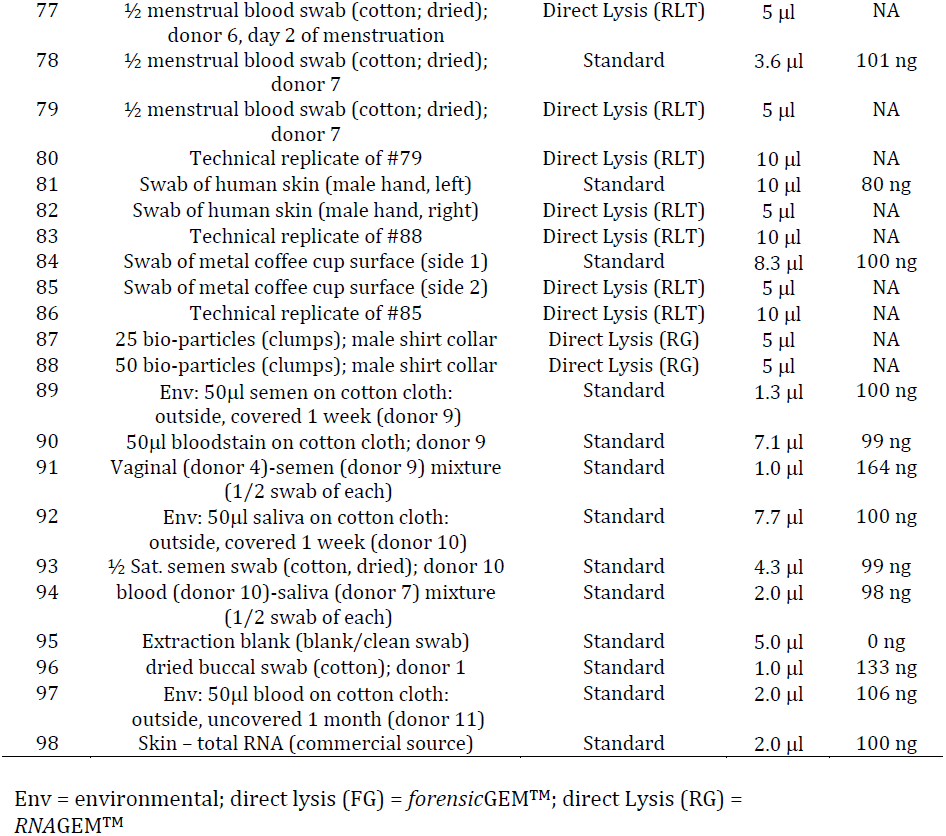
Sample Descriptions and Assay Input (Full Sample Set)

**Supplementary Figure 1.**
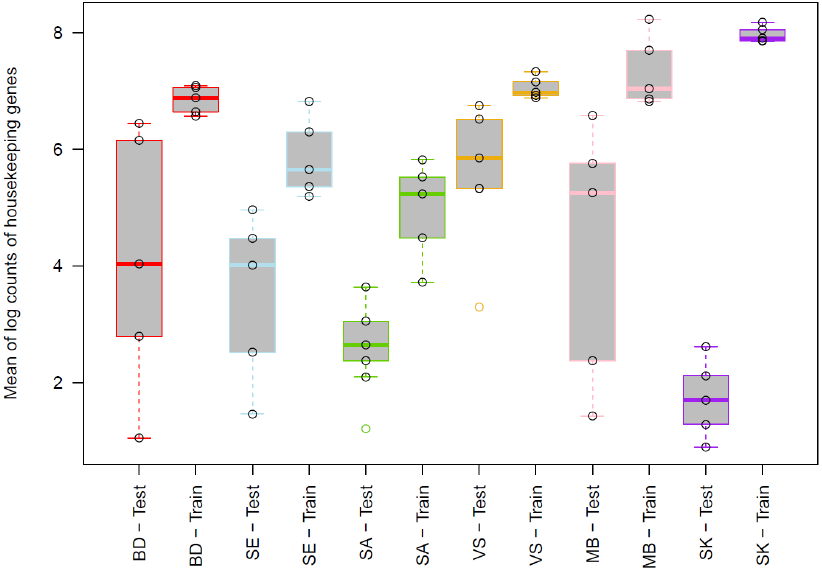
Housekeeping gene expression in training and test samples.

**Supplementary Figure 2.**
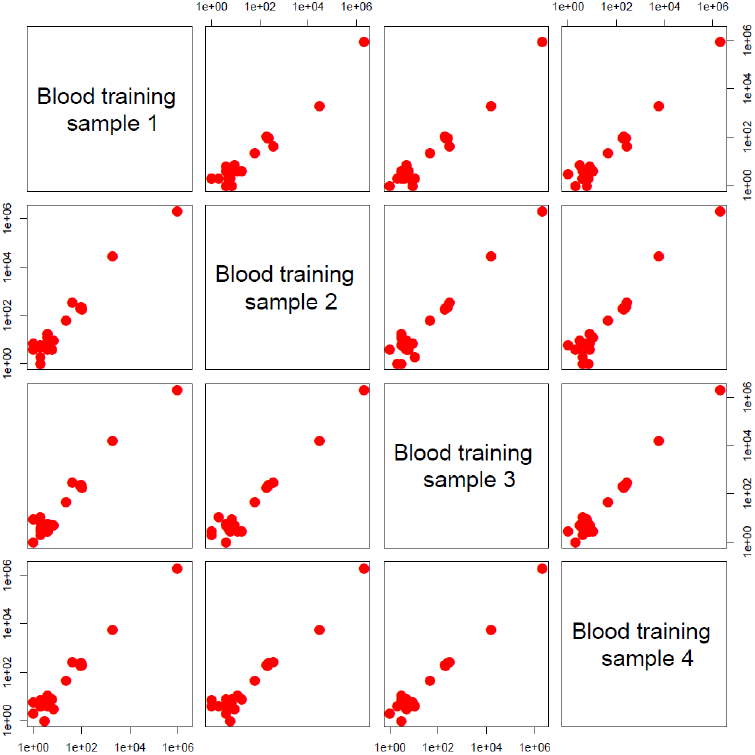
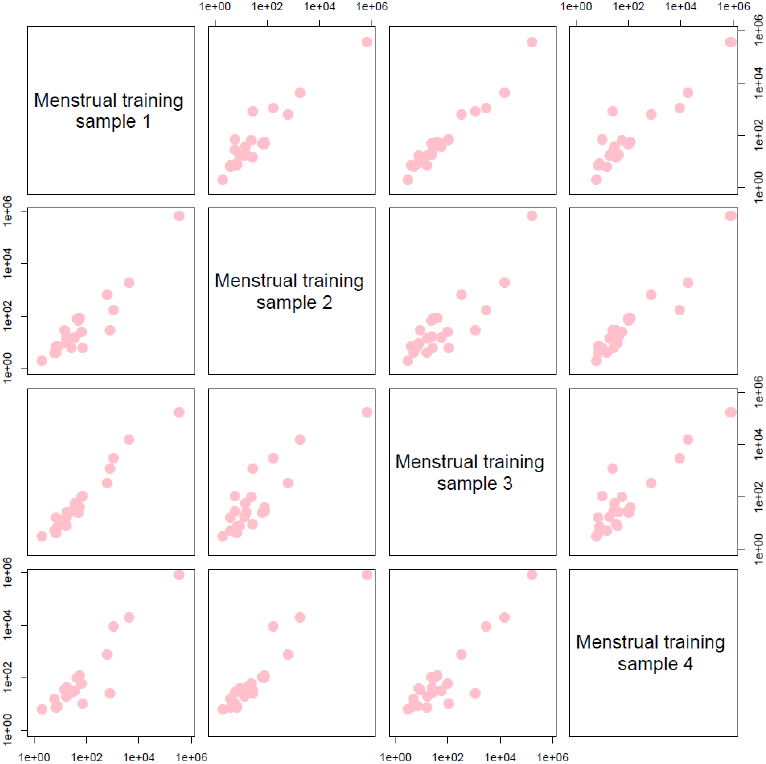
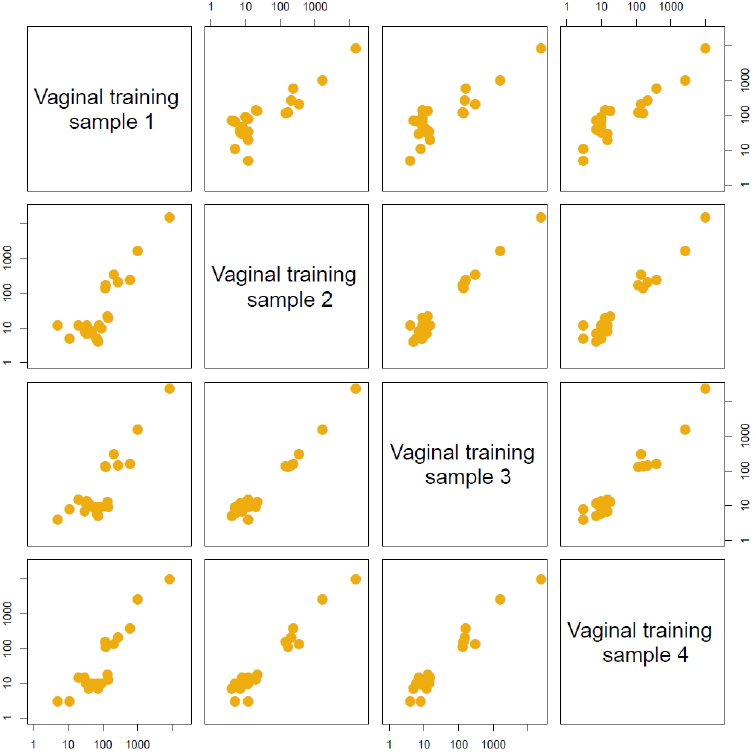
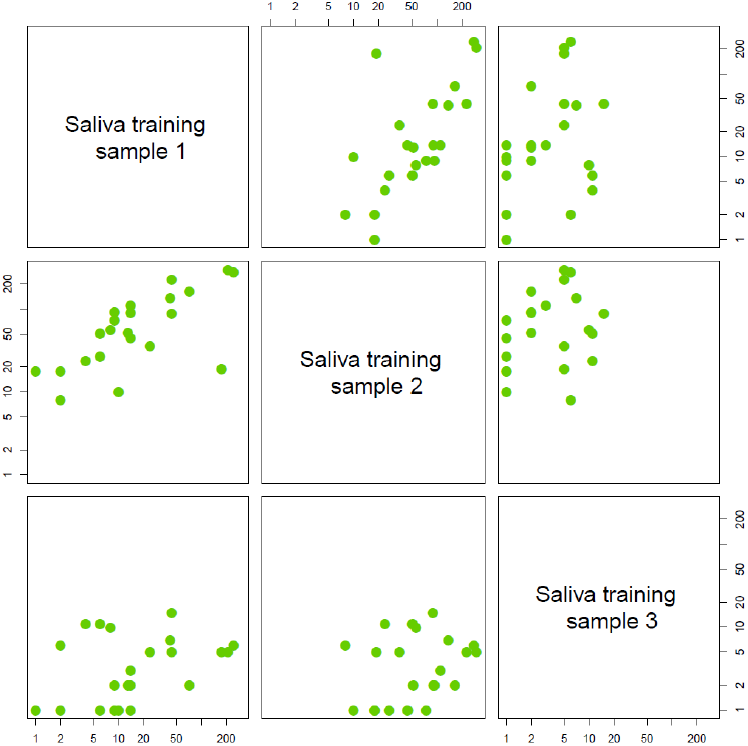
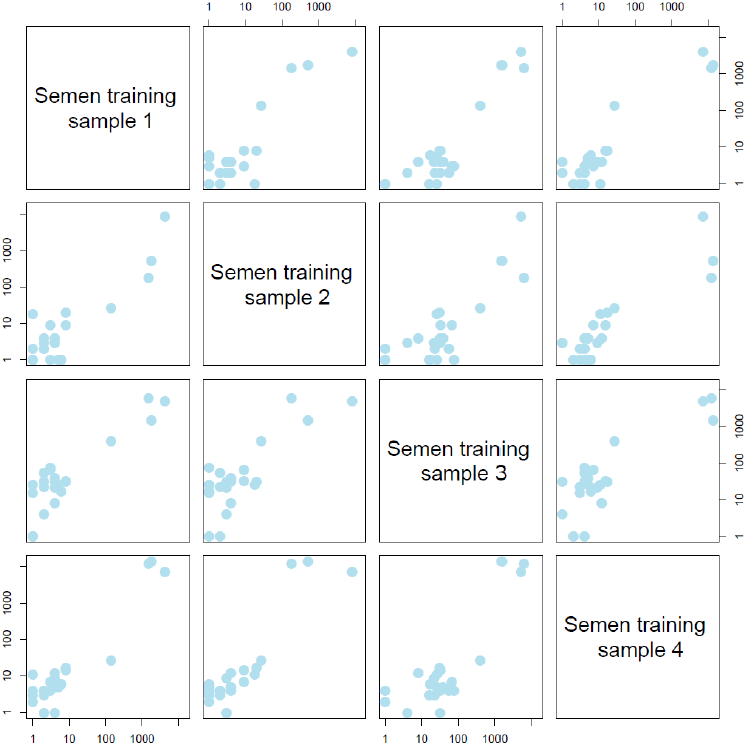
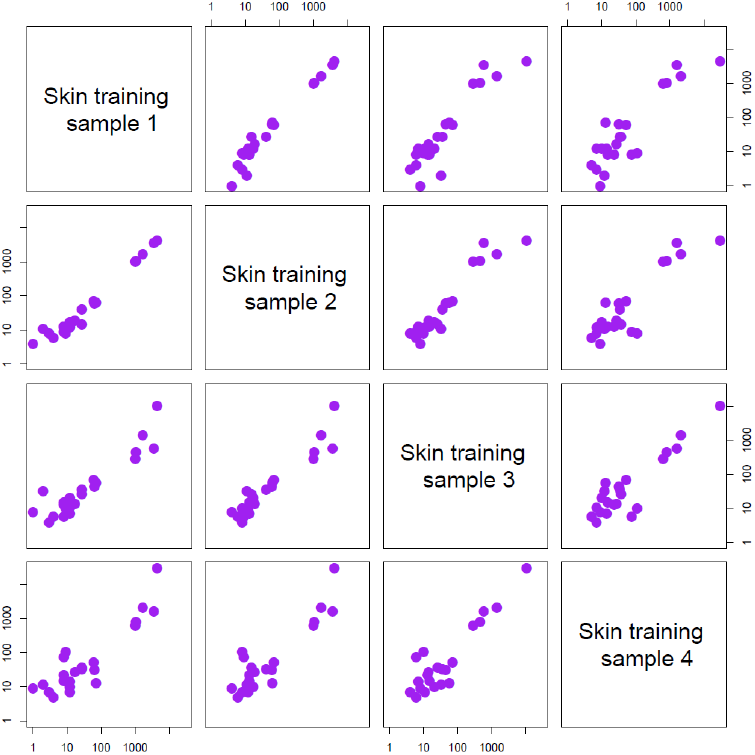
Profiles of the training samples from each fluid are plotted against each other.

**Supplemental Figure 3.**
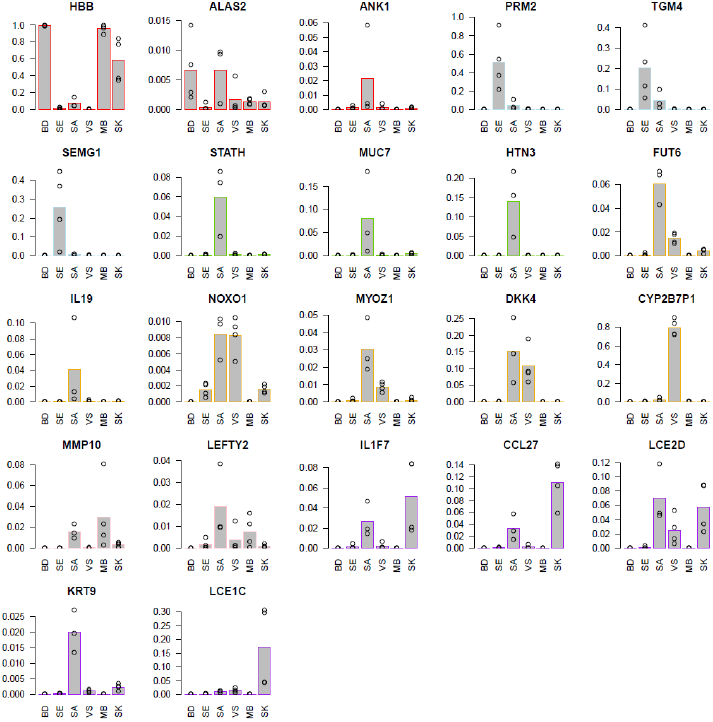
Boxplots for individual genes’ proportion of total expression in the different sample types.

BD = blood, SE =semen, SA = saliva, MB = menstrual blood, VS = vaginal secretions, SK = skin.

**Supplementary Figure 4.**
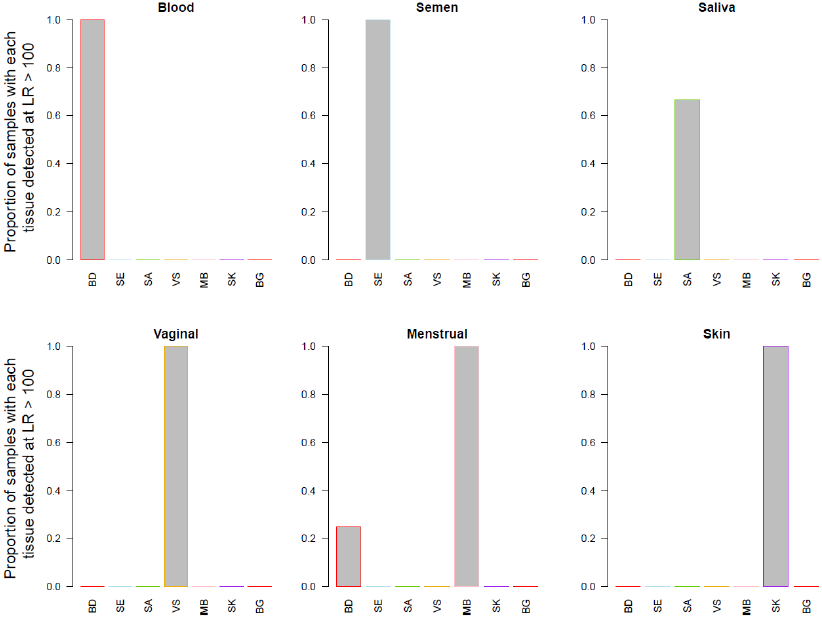
Performance of the algorithm in the training set.

Bars display the rate at which each fluid is called detected in each sample type. Fluids are called detected if their likelihood ratio exceeds 100.

**Supplementary Figure 5.**
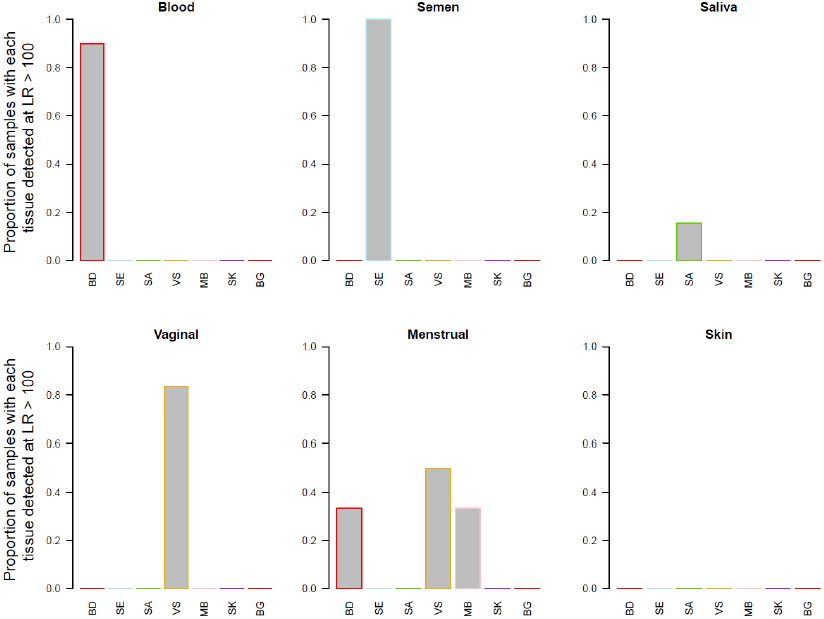
Performance of the algorithm in the test set.

Bars display the rate at which each fluid is called detected in each sample type. Fluids are called detected if their likelihood ratio exceeds 100.

**Supplementary Figure 6.**
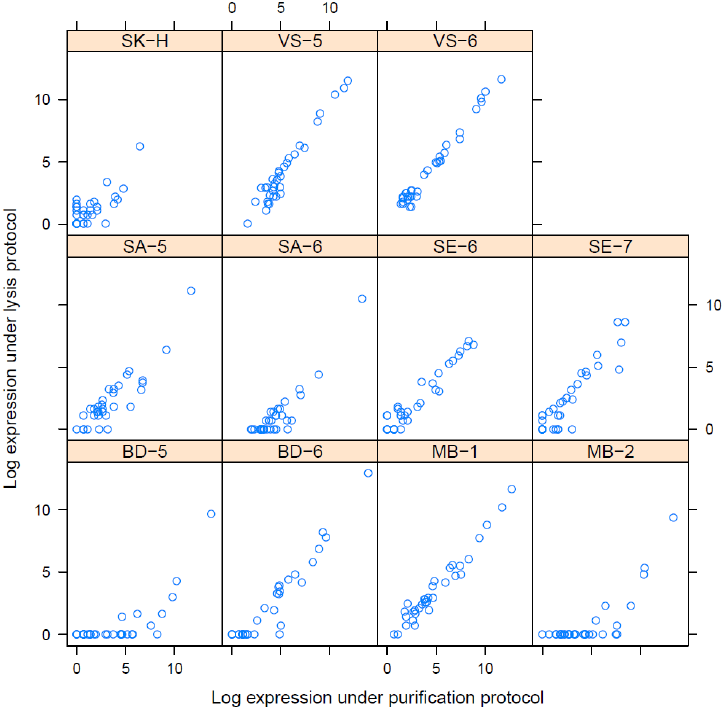
Concordance of the assay between purification and lysis protocols.

For the 14 samples with replicates run under each protocol, the natural log gene expression profile under the lysis protocol (vertical axis) is plotted against the profile under the purification protocol (horizontal axis).

